# Large-scale epidemiological study on feline autosomal dominant polycystic kidney disease and identification of novel *PKD1* gene variants

**DOI:** 10.1101/2023.03.23.533873

**Authors:** Fumitaka Shitamori, Ayaka Nonogaki, Tomoki Motegi, Yuki Matsumoto, Mika Sakamoto, Yasuhiro Tanizawa, Yasukazu Nakamura, Tomohiro Yonezawa, Yasuyuki Momoi, Shingo Maeda

**Affiliations:** Department of Veterinary Clinical Pathobiology, Graduate School of Agricultural and Life Sciences, The University of Tokyo, Tokyo, Japan; Veterinary Medical Center, The University of Tokyo, Tokyo, Japan; Anicom Specialty Medical Institute Inc., Kanagawa, Japan; Department of Informatics, National Institute of Genetics, Research Organization of Information and Systems, Shizuoka, Japan

**Keywords:** ADPKD, cat, epidemiology, genotyping, target resequencing

## Abstract

Autosomal dominant polycystic kidney disease (ADPKD) is a common inherited disease in cats. In most cases, the responsible abnormality is a nonsense single nucleotide polymorphism in exon 29 of the *PKD1* gene (chrE3:g.42858112C>A, the conventional *PKD1* variant). Epidemiological studies on feline ADPKD caused by the conventional *PKD1* variant have been conducted in several countries, including Japan. However, they were limited to Persian cats or cats already suspected of ADPKD. There are no data on the prevalence of the conventional *PKD1* variant among a wider cat population in Japan that is not limited to certain breeds or clinical backgrounds. This large-scale epidemiological study involving 1,281 cats aimed to establish a new genotyping assay to detect the conventional *PKD1* variant rapidly. Among the cats examined in this study, 1.8% (23/1,281) harbored the conventional *PKD1* variant. The odds of having the conventional *PKD1* variant were significantly higher in Persian cats, Scottish Folds, and Exotic Shorthairs than in the other breeds. Furthermore, targeted resequencing of *PKD1* was performed in cats with ADPKD and healthy cats to search for new variants that may be involved in ADPKD. We identified four variants unique to ADPKD cats that were not found in healthy cats, all of which were in exon 15. This indicates that variants in exon 15 of *PKD1*, in addition to the conventional variant in exon 29, are key factors in the pathogenesis of ADPKD in cats.

## INTRODUCTION

Autosomal dominant polycystic kidney disease (ADPKD) is a common inherited disease in cats and is characterized by the development of multiple fluid-filled cysts dispersed in the kidney and occasionally in the liver or pancreas [12]. The cysts gradually increase in number and size, causing deterioration of the renal function through compression of the renal parenchyma. Most patients remain clinically normal for years until chronic kidney disease (CKD) develops later in life (>7 years) [7]. Ultrasonographic examination is routinely performed for the diagnosis and evaluation of feline ADPKD. Owing to its hereditary nature, no fundamental treatment options preventing the onset of the disease have been found to date. Hence, feline ADPKD is generally treated by following the standard CKD treatment protocols such as subcutaneous fluid administration [27].

Feline ADPKD is caused by a heterozygous variant that was identified by Lyons et al. in 2004 and is located in the *PKD1* gene with a nonsense single nucleotide polymorphism (SNP) in exon 29 (chrE3:g.42858112C>A, the conventional *PKD1* variant). This variant causes a stop codon, truncating the protein up to 25% at the C-terminus [21]. This variant in a homozygous state is lethal in the embryonic stage. *PKD1* encodes polycystin 1 (PC1), which is a membrane-bound large multidomain glycoprotein with a G-protein-coupled receptor proteolytic site. Its physiological function is unclear; however, it is known to be located on the primary cilia, a flagellar organelle protruding from the cell surface used for mechanotransduction. PC1 is thought to perform an important role in transmitting signals from outside the cell [10, 15]. The loss of functional PC1 caused by the variant appears to contribute to the cystogenesis in ADPKD, and much research has been conducted to identify the underlying mechanisms. Some have proposed the influence of various upregulated or downregulated intracellular signals, while others have argued that macrophage recruitment by the primary cilia plays a huge role in cystogenesis [16]. Despite these efforts, the exact mechanism remains unclear. However, two models on the relationship between PC1 dosage and cystogenesis have been established: the two-hit model and the threshold model. The two-hit model suggests that after inactivation of one allele by germline mutation, a somatic mutation occurs, inactivating the other allele and thereby causing cystogenesis [24]. On the other hand, the threshold theory is the more recent belief, which suggests that the disease occurs when the PC1 levels drop below a certain threshold and do not necessarily have to be at zero [14, 17]. Since the variant responsible for feline ADPKD truncates PC1, it can be assumed that the truncated abnormal protein is unable to function properly, leading to a lack of functional PC1. Subsequent to the discovery of the responsible variant, *PKD1* genetic tests have also been developed for the diagnosis of feline ADPKD, which allows for easier diagnosis even in the asymptomatic early stages of ADPKD [29]. However, these tests cannot predict the severity of the disease or evaluate its progression, suggesting that genetic tests should be used together with ultrasonographic examination.

Previous studies in the United States, Europe, Australia, and Taiwan have suggested that ADPKD in cats is most common in Persian cats and related breeds [2, 3, 8, 12, 18]; the same trend has been reported in Japan along with reports of ADPKD in other breeds such as Scottish Folds and American Shorthairs [25]. The prevalence of the conventional *PKD1* variant has been investigated in very few studies and reported as 13.5% in China (n=111), 33.3% in Slovenia (n=24, limited to Persian cats), and 40% in Japan (n=377, limited to cats suspected of ADPKD) [11, 18, 25]. These studies have mostly focused on Persian cats or cats already suspected of ADPKD. There are no data on the prevalence of the disease or the conventional *PKD1* variant among the cat population as a whole, even though comprehending the frequency of the conventional *PKD1* variant and confirming predisposed breeds in every region is crucial for prevention of ADPKD. Additionally, some studies have revealed that cats without the conventional *PKD1* variant can develop ADPKD. Three cases of ADPKD diagnosis based on ultrasonographic examination in the study in China did not have the conventional *PKD1* variant [18]. Another study on Maine Coons in Switzerland reported that none of the six cases with ADPKD had the *PKD1* conventional variant [13]. These observations suggest that an unknown variant may serve as another cause of ADPKD.

Given this background of feline ADPKD, we conducted a large-scale epidemiological research to understand the nature and current situation of ADPKD in Japan accurately. Additionally, we performed target resequencing of all exons of *PKD1* using next-generation sequencing to identify new causative variants.

## MATERIALS AND METHODS

### Feline cases and samples

There were 7,536 visits and 5,716 admissions of feline cases at the Veterinary Medical Center of the University of Tokyo between April 2015 and March 2018 (cumulative total, 13,252). After excluding repetitive visits and patients that did not undergo blood tests, 1,281 cats were included in this study.

Clinical information on the age, breed, sex, and diagnosis of all cases was obtained from the electronic medical records system. The cats were 0–20 years of age, comprising 717 males (101 intact and 616 neutered), 560 females (94 intact and 466 neutered), and 4 of unknown sex. There was a wide variety of breeds as follows: mixed (n=791), American Shorthair (n=101), Scottish Fold (n=87), Persian cat (n=37), Russian Blue (n=34), Maine Coon (n=31), Norwegian Forest cat (n=29), Munchkin (n=28), Abyssinian (n=24), Somali (n=20), Ragdoll (n=16), British Shorthair (n=14), American Curl (n=13), Himalayan (n=13), Bengal (n=13), Chartreux (n=6), Exotic Shorthair (n=6), Singapura (n=6), Ocicat (n=2), Ragamuffin (n=2), Siamese (n=2), Tonkinese (n=2), Birman (n=1), European Shorthair (n=1), La Perm (n=1), and Selkirk Rex (n=1). The body weight, ultrasonographic findings, and values of blood urea nitrogen (BUN), creatinine, and urine specific gravity were assessed of the cats with the conventional *PKD1* variant according to genotyping from the electronic medical record system. The electronic medical records system was scanned for the mention of feline ADPKD in the same period (April 2015 to March 2018) to identify samples diagnosed as ADPKD that did not have the conventional *PKD1* variant according to genotyping. Ultrasonographic images were retrieved and reviewed for the applicable samples.

The Animal Care and Clinical Research Committees of the Veterinary Medical Center of the University of Tokyo approved the study protocols for blood sampling and collection of clinical information of the client-owned cats. DNA was extracted from ethylenediaminetetraacetic acid (EDTA)-anticoagulated blood using NucleoSpin Blood Quick Pure kit (Macherey-Nagel, Duren, Germany) and was stored in the gene bank run by Azabu University (Kanagawa, Japan).

### Real-time polymerase chain reaction PKD1 genotyping

We conducted a real-time polymerase chain reaction (PCR) genotyping assay using TaqMan probes, which are probes with fluorescein amidite (FAM) and 2’-chloro-7’phenyl-1,4-dichloro-6-carboxy-fluorescein (VIC) on one end and a nonfluorescent quencher (NFQ) on the other. The assay was performed with 5.0 μL TaqPath ProAmp Master Mix (Thermo Fisher Scientific, Waltham, MA, USA), 0.5 μL Custom TaqMan SNP Genotyping Assay (Thermo Fisher Scientific, Waltham, MA, USA), >20 ng DNA, and ultrapure water up to 10 μL. The Custom TaqMan SNP Genotyping Assay included forward (CCTCGGAGCCGCTTCAC) and reverse (TTGGCGCCCAGGAAGAG) primers and two probes, VIC–ACGAGGAGGACGCAACA–NFQ and FAM–CGAGGAGGACTCAACA–NFQ, for wild-type and mutant alleles, respectively. Real-time PCR was performed on the fast cycle on StepOnePlus Real-time PCR system (Thermo Fisher Scientific, Waltham, MA, USA) for 1 cycle at 60°C for 30 sec and 95°C for 5 min. This was followed by 40 cycles of 95°C for 5 sec and 60°C for 30 sec and then 1 cycle of 60°C for 30 sec in the end.

To determine the specificity of this assay, two samples each of wild-type and heterozygous mutant samples selected randomly and a water control were run on the genotyping assay. The genotypes of these samples were confirmed by Sanger sequencing (Azenta Life Sciences, Tokyo, Japan) after PCR amplification and extrication from the agarose gel using Wizard^®^ SV Gel and PCR Clean-Up System (Promega, Madison, WI, USA). The threshold was set to 0.6 ΔRn, and threshold cycle values were calculated automatically on the StepOnePlus Real-time PCR system. A value >40 was assigned to samples whose threshold cycle values were undetermined after 40 PCR cycles.

Real-time PCR assay was performed as described above for all DNA feline samples to determine their genotype. The genotypes of each sample were determined by examining the amplification of the reporter dye (FAM or VIC) on the amplification plot.

### Target resequencing of PKD1 using next-generation sequencing

This study included 23 feline cases with renal cysts confirmed through ultrasonographical examination and also with the conventional *PKD1* variant (chrE3:g.42858112C>A), 6 cases of ADPKD with renal cysts but without the conventional *PKD1* variant, and 62 cases without renal cysts and the conventional *PKD1* variant. DNA was extracted from EDTA-anticoagulated blood using NucleoSpin Blood Quick Pure kit and was stored with help from the gene bank run by Azabu University. Data regarding age, breed, sex, diagnosis, and ultrasonographic images of all patients were obtained from the electronic medical records system.

We designed two primer pools of 106 primer pairs with DesignStudio (Illumina, San Diego, CA, USA) to amplify *PKD1*. The sequencing library was constructed using the AmpliSeq Library PLUS for Illumina. The PCR product size distribution was checked using Bioanalyzer (Agilent Technologies, Santa Clara, CA, USA) after adapter ligation. Paired-end sequencing (300 bp) with a dual 8-bp barcode was performed using MiSeq (Illumina, San Diego, CA, USA). The sequence reads were sorted individually using each barcode.

Raw sequence data were processed according to the Broad Institute’s Genome Analysis Tool Kit (GATK) best practices workflow for small germline variants [28]. Raw sequence reads were trimmed to remove adaptors using Trimmomatic (version 0.38) [6] and mapped to the feline reference sequence (Ensembl Genome assembly felCat9 GCA_000181335.4) using bwa mem (version 0.7.17) [19]. Samtools (version 1.9) was used to sort and index each mapped file [20]. These data were subjected to VariantFiltration using GATK (version 4.2.2). Moreover, the data were applied to VariantFiltration using GATK (version 4.2.2) with the filter-expression parameter of “QD < 5.0 || FS > 50.0 || SOR > 3.0 || MQ < 50.0 || MQRankSum < –2.5 || ReadPosRankSum < –1.0 || ReadPosRankSum > 3.5.” In individual samples, the coverage rate was defined as the proportion of the target region covered by ≥20 sequencing reads. Variants were called using HaplotypeCaller by GATK. Annotation was performed using SnpEff (version 4.3) [9] and Ensembl Variant Effect Predictor [23], with all transcripts registered in Ensembl release 104.

### Statistical analyses

Statistical analyses were performed using GraphPad Prism software (version 5.0.1; GraphPad Software, San Diego, CA, USA). The odds ratio was calculated for each breed to determine the breeds predisposed to the conventional *PKD1* variant. Significance was confirmed using two-tailed Fisher exact test. The relationship between the frequency of the conventional *PKD1* variant and sex was evaluated using the chi-square test. The relationship between the presence of renal cysts and the age and plasma creatinine or BUN levels was determined using two-tailed Mann–Whitney U test. *P*-values <0.05 were considered statistically significant.

## RESULTS

### Establishment of a novel TaqMan real-time PCR genotyping assay

To determine the specificity of the real-time PCR *PKD1* genotyping assay, two samples each of heterozygous *PKD1* mutants and wild-type cases, whose genotypes had previously been confirmed by Sanger sequencing, were randomly selected. The two wild-type samples produced threshold cycle values in the VIC channel of 29.165 and 28.124, respectively. However, these samples did not produce threshold cycle values within 40 cycles in the FAM channel (>40 cycles were needed) (Table 1). On the other hand, the two heterozygous mutant samples had similar threshold values in both the VIC (29.564 and 29.957, respectively) and FAM channels (28.041 and 28.684, respectively) (Table 1). Figure 1 shows the production of threshold cycle values for water, *PKD1* wild-type, and *PKD1* variant samples. Wild-type samples showed a slight increase in the ΔRn in FAM, which could be considered non-specific (Fig. 1B). This increase was within the threshold range (ΔRn=0.6), even within 40 cycles, and was clearly different from the increase in FAM in *PKD1* mutants (Fig. 1C). These results suggest that the assay was able to discriminate between wild-type and the conventional *PKD1* variant with high specificity.

**Table 1.**
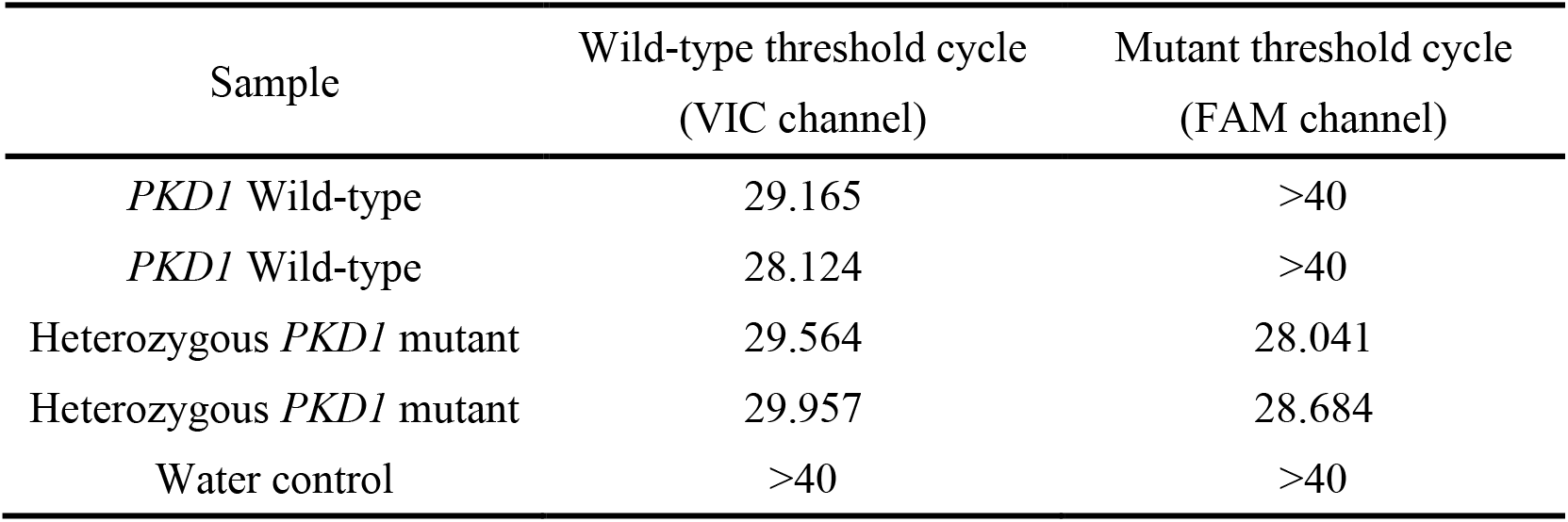
Specificity of the real-time PCR genotyping assay

**Fig. 1.**
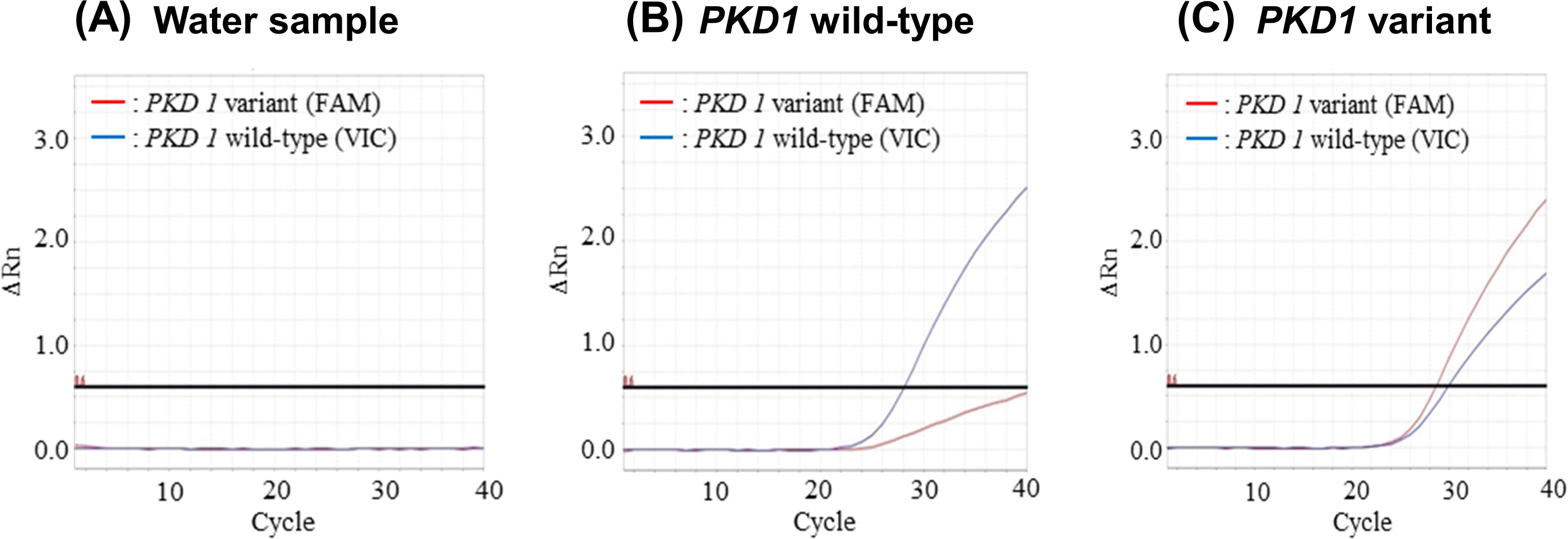
Representative amplification plots from real-time polymerase chain reaction genotyping The red graph shows amplification of the reporter dye on the conventional *PKD1* variant probe (FAM) and the blue graph of the wild probe (VIC). (**A**) In the water sample, no changes were noted in either graph. (**B**) In the wild-type samples, only the blue graph showed a significant increase. The slight increase in the red graph is presumably nonspecific. (**C**) In the DNA samples with the conventional *PKD1* variant (chrE3:g.42858112C>A), both graphs showed increased values, reflecting the heterozygous variant. (FAM, fluorescein amidite; VIC, 2’-chloro-7’phenyl-1,4-dichloro-6-carboxy-fluorescein)

### Prevalence of the conventional PKD1 variant and clinical evaluation

Out of 1,281 samples, 23 (1.8%) had the conventional *PKD1* variant (chrE3:g.42858112C>A), while 9 diagnosed with ADPKD did not have the conventional *PKD1* variant (Table 2). The 23 samples comprised Scottish Folds (n=6), mixed breed (n=6), Persian cats (n=4), American Shorthairs (n=3), Exotic Shorthairs (n=2), and Munchkins (n=2). The breed with the highest positive rate for the variant was Exotic Shorthair at 33.3%. Persian cats also had a high positive rate of 10.8%, followed by Munchkin (7.14%), Scottish Fold (6.90%), and American Shorthair (2.97%) breeds (Table 3). The odds of having the conventional *PKD1* variant were significantly higher in Persian cats, Scottish Folds, and Exotic Shorthairs than in the other breeds (Table 3). Cases with the conventional *PKD1* variant were dispersed across different age and sex groups, with an age range of 0–13 years and comprising 14 females (three intact and 11 neutered) and 9 males (all neutered). There was no significant difference in the frequency of the conventional *PKD1* variant according to the sex by the chi-square test (*P*=0.0988).

**Table 2.** Cases with the *PKD1* variant confirmed by genotyping in this study and cases diagnosed with ADPKD without the *PKD1* variant

| Cases with the *PKDI* variant |  |  |  |  |
| --- | --- | --- | --- | --- |
| Case | Years | Sex<sup>*1</sup> | Breed | Renal cysts<sup>*2</sup> |
| 1 | 5 | SF | American Shorthair | + |
| 2 | 5 | SF | Persian | + |
| 3 | 5 | SF | Mixed | + |
| 4 | 5 | CM | Munchkin | + |
| 5 | 1 | SF | Scottish Fold | NE |
| 6 | 9 | CM | Scottish Fold | + |
| 7 | 0 | SF | Scottish Fold | - |
| 8 | 13 | CM | Mixed | + |
| 9 | 8 | Female | Mixed | Pancreatic |
| 10 | 5 | CM | Persian | Liver |
| 11 | 0 | Female | Mixed | - |
| 12 | 9 | CM | Persian | + |
| 13 | 2 | CM | Mixed | + |
| 14 | 5 | SF | Mixed | NE |
| 15 | 3 | SF | Scottish Fold | + |
| 16 | 10 | SF | Exotic Shorthair | + |
| 17 | 9 | SF | Scottish Fold | + |
| 18 | 1 | SF | Scottish Fold | + |
| 19 | 9 | CM | Exotic Shorthair | + |
| 20 | 12 | CM | American Shorthair | + |
| 21 | 9 | CM | Munchkin | + |
| 22 | 9 | Female | Persian | + |
| 23 | 1 | SF | American Shorthair | NE |

| ADPKD without the *PKD1* variant |  |  |  |  |
| --- | --- | --- | --- | --- |
| Case | Years | Sex | Breed | Renal cysts |
| 1 | 13 | SF | Mixed | + |
| 2 | 14 | CM | American Shorthair | + |
| 3 | 13 | SF | Mixed | + |
| 4 | 10 | CM | Singapura | + |
| 5 | 14 | CM | Mixed | + |
| 6 | 12 | SF | Scottish Fold | + |
| 7 | 9 | Female | American Shorthair | + |
| 8 | 1 | CM | Scottish Fold | + |
| 9 | 10 | SF | Scottish Fold | + |

\*<sup>1</sup> SF=spayed female; CM=castrated male
\*<sup>2</sup> NE=not evalulated; Pancreatic=pancreatic cyst only; Liver=liver cyst only

**Table 3.** Prevalence of the PKD1 variant and odds in different breeds

| Breed | *PKDI*<br>variants | Wild-type | Prevalence<br>(%) | Odds ratio | 95% Confidence<br>Interval | *P*-value |
| --- | --- | --- | --- | --- | --- | --- |
| Scottish Fold | 6 | 81 | 6.89 | 5.129 | 1.968-13.36 | **0.0033** |
| Persian | 4 | 33 | 10.81 | 7.815 | 2.518-24.25 | **0.0035** |
| American Shorthair | 3 | 98 | 2.97 | 1.776 | 0.5184-6.082 | 0.4187 |
| Exotic Shorthair | 2 | 4 | 33.33 | 29.86 | 5.179-172.1 | **0.0044** |
| Munchkin | 2 | 26 | 7.14 | 4.513 | 1.005-20.26 | 0.0879 |

Renal cysts were confirmed on ultrasonography in 16 of the 23 cats with the conventional *PKD1* variant, 4 cats did not have renal cysts, and the remaining 3 cats did not undergo ultrasonography (Table 2). There was no statistically significant relationship between age and the presence of renal cysts; however, there was a higher tendency for renal cysts occurrence in older cats than in the younger cats (*P*=0.0598; Fig. 2). Two out of the four cats without renal cysts were under 1 year of age, while the other two were relatively older (5 and 8 years, respectively); one of the older cats had multiple cysts in the liver and the other in the pancreas.

**Fig. 2.**
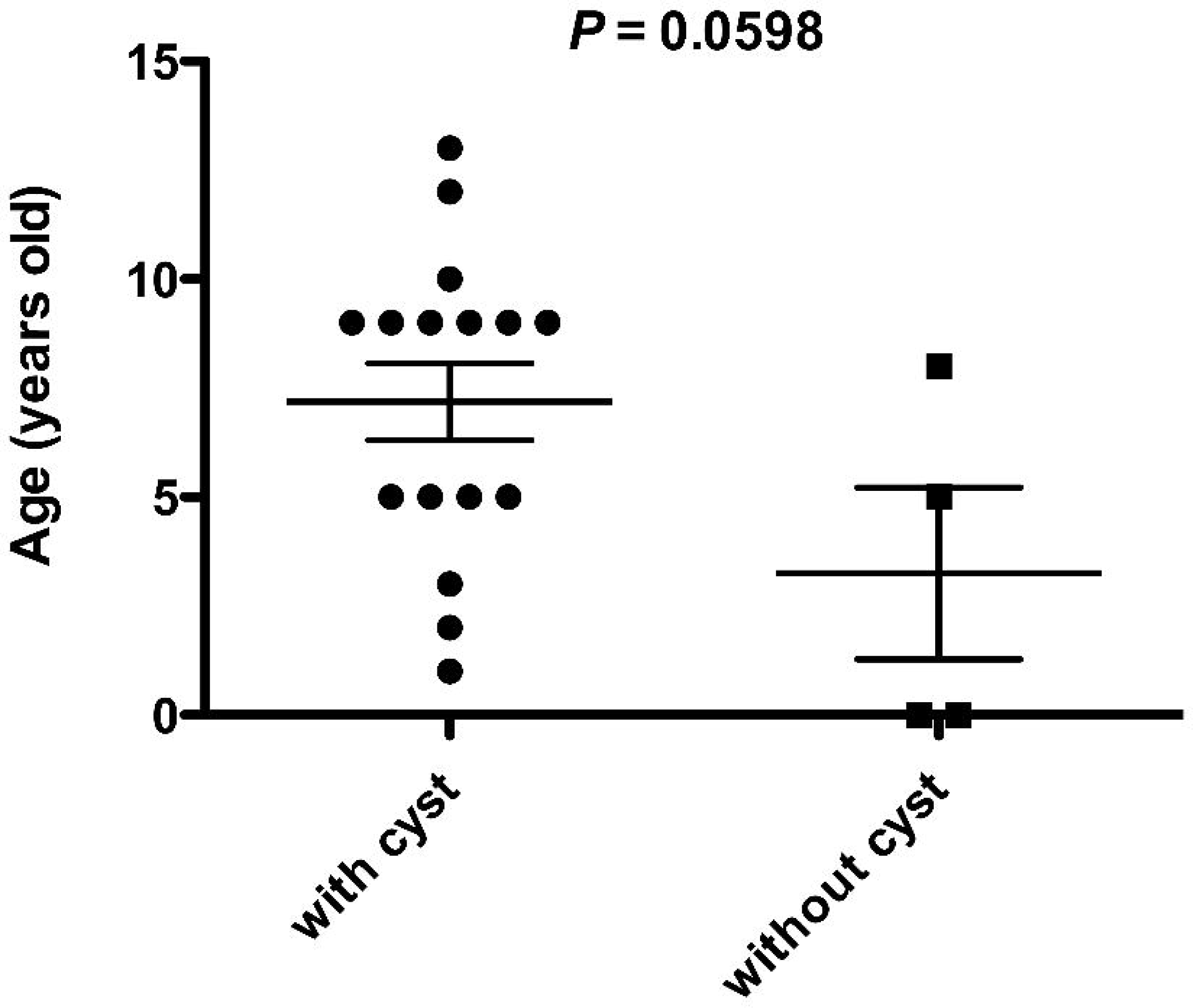
Association between the age and presence of renal cysts in cats with the conventional *PKD1* variants (n=20) Of the 23 cases with the conventional *PKD1* variants, 3 cases did not undergo ultrasound examination. Results are expressed as means ± standard errors. Data were evaluated using two-tailed Mann–Whitney U test (*P*=0.0598).

Plasma creatinine and BUN levels were compared between the patients with and without renal cysts (Fig. 3). BUN levels were significantly higher in cats with renal cysts than in those without (*P*=0.0178), and even plasma creatinine levels tended to be higher in the former (*P*=0.0992).

**Fig. 3.**
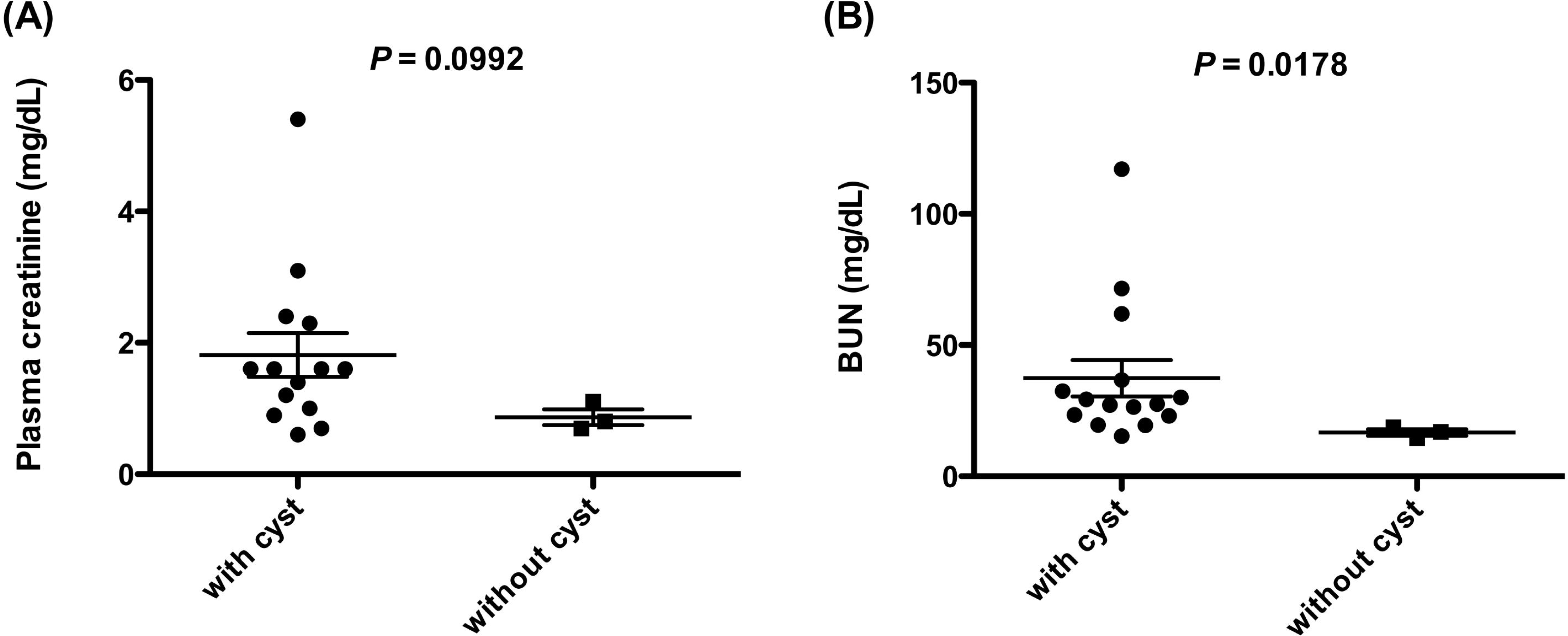
Association between presence of renal cysts and the (**A**) plasma creatinine (n=17) and (**B**) blood urea nitrogen (n=18) levels in cats with the conventional *PKD1* variants Of the 23 cases with the conventional *PKD1* variants, 3 cases did not undergo ultrasound examination. Additionally, the plasma creatinine values were not recorded in three cases, while BUN values were not recorded in two; hence, they were excluded from this analysis. Results are expressed as means ± standard errors. Data were evaluated using two-tailed Mann–Whitney U test (plasma creatinine: *P*=0.0992, BUN: *P*=0.0178). (BUN, blood urea nitrogen)

### Target resequencing of PKD1 using next-generation sequencing

One of 62 controls and one of 23 ADPKD cats with the conventional *PKD1* variant (chrE3:g.42858112C>A) were excluded because their DNA concentrations were low. We identified 86 nonsynonymous variants of *PKD1*, including 18 protein-truncating variants and 68 missense variants, among the included samples. Of these variants, 27 nonsynonymous variants (2 protein-truncating variants and 25 missense variants) were identified in six cases of ADPKD without the conventional *PKD1* variant (Table 4). We focused on four singleton variants that were found only in six cases of ADPKD without the conventional *PKD1* variant but not in any of the control cases (variants #4, #9, #14, and #18). All four of these variants were located in exon 15, of which one was a single-base deletion (chrE3:g.42848725delC), which causes immense damage to the resulting protein, and the other three were missense variants (Table 5). The three missense variants were analyzed using SIFT [26] and PolyPhen-2 [1] tools to estimate their impact on amino acid substitutions. We found that chrE3:g.42848361A>C (p.His1520Pro) and chrE3:g.42849470G>C (p.Val1890Leu) were predicted as benign in both software packages. However, chrE3:g.42850283C>T (p.Arg2162Trp) was predicted as damaging by PolyPhen-2 (Table 5). Variants chrE3:g.42848361A>C and chrE3:g.42849470G>C were present in the same cat. We used the UCSC LiftOver tool (https://genome.ucsc.edu/cgi-bin/hgLiftOver) to compare the variant locations with the human gene variants reported in human ADPKD. We found that chrE3:g.42850283C>T was identical to the human variant rs373256534 (p.Arg2162Trp).

**Table 4.** Novel *PKD1* gene variants found in ADPKD cases without the chrE3:g.42858112C>A nonsense variant

| Variant No. | Location | ADPKD<br>(n=28) |  | Healthy<br>(n=61) |  |
| --- | --- | --- | --- | --- | --- |
|  |  | + | – | + | – |
| 1 | chrE3:g.42843634A>G | 25 | 3 | 34 | 27 |
| 2 | chrE3:g.42844184C>A | 25 | 3 | 36 | 25 |
| 3 | chrE3:g.42846440A>G | 5 | 23 | 5 | 56 |
| 4 | chrE3:g.42847359A>C | 25 | 3 | 34 | 27 |
| 5 | chrE3:g.42847502A>G | 25 | 3 | 39 | 22 |
| 6 | chrE3:g.42848285A>T | 25 | 3 | 33 | 28 |
| 7 | **chrE3:g.42848361A>C** | **1** | 27 | **0** | 61 |
| 8 | chrE3:g.42848369A>G | 25 | 3 | 52 | 9 |
| 9 | **chrE3:g.42848725delC** | **1** | 27 | **0** | 61 |
| 10 | chrE3:g.42848883G>A | 7 | 21 | 35 | 26 |
| 11 | chrE3:g.42849191G>T | 2 | 26 | 4 | 57 |
| 12 | chrE3:g.42849230G>A | 2 | 26 | 4 | 57 |
| 13 | chrE3:g.42849461G>A | 25 | 3 | 36 | 25 |
| 14 | **chrE3:g.42849470G>C** | **1** | 27 | **0** | 61 |
| 15 | chrE3:g.42849546C>G | 9 | 19 | 31 | 30 |
| 16 | chrE3:g.42849762T>C | 28 | 0 | 58 | 3 |
| 17 | chrE3:g.42850022C>T | 2 | 26 | 1 | 60 |
| 18 | **chrE3:g.42850283C>T** | **1** | 27 | **0** | 61 |
| 19 | chrE3:g.42850889A>G | 25 | 3 | 36 | 25 |
| 20 | chrE3:g.42853138A>G | 25 | 3 | 36 | 25 |
| 21 | chrE3:g.42853493A>G | 25 | 3 | 34 | 27 |
| 22 | chrE3:g.42853544A>G | 25 | 3 | 34 | 27 |
| 23 | chrE3:g.42855067G>C | 27 | 1 | 53 | 8 |
| 24 | chrE3:g.42862845A>G | 28 | 0 | 60 | 1 |
| 25 | chrE3:g.42864749C>T | 25 | 3 | 35 | 26 |
| 26 | chrE3:g.42864765T>C | 28 | 0 | 54 | 7 |
| 27 | chrE3:g.42866743delC | 2 | 26 | 14 | 47 |

**Table 5.** Impact of *PKD1* variants estimated by SIFT and PolyPhen-2 software

| Variant No. | Location | Consequence | Exon | cDNA position | Impact analysis |  |
| --- | --- | --- | --- | --- | --- | --- |
|  |  |  |  |  | SIFT | Poly-phen2 |
| 7 | chrE3:g.42848361A>C | missense<br>(p.His1520Pro) | 15 | 4886 | tolerated<br>(0.28) | benign<br>(0.170) |
| 9 | chrE3:g.42848725delC | frameshift<br>(p.Gly1641fs) | 15 | 5250 | - | - |
| 14 | chrE3:g.42849470G>C | missense<br>(p.Val1890Leu) | 15 | 5995 | tolerated<br>(0.11) | benign<br>(0.103) |
| 18 | chrE3:g.42850283C>T | missense<br>(p.Arg2162Trp) | 15 | 6808 | tolerated<br>(0.28) | probably<br>damaging<br>(0.999) |

## DISCUSSION

Feline ADPKD first featured in case reports in the 1960s as a disease observed in Persian and Persian-cross cats. Subsequent research on the disease has been based on this presupposition, mainly focusing on Persian cat-related breeds [4, 5]. Moreover, the conventional *PKD1* variant (chrE3:g.42858112C>A) was initially discovered in Persian cats [21]. Occurrence of the disease in other breeds has also been reported more recently [18], shedding light on the importance of investigating the disease on a wider scale to capture the whole picture. Although a recent study explored the prevalence of the conventional *PKD1* variant in Japan (n=377), it had limitations in that it focused only on cases suspected of ADPKD based on family history or ultrasonographic observations [25]. In contrast, this study investigated 1,281 cats, regardless of the case history, chief complaint, diagnosis, or breed, making it the first large-scale epidemiological study assessing the conventional *PKD1* variant prevalence in Japan using a novel real-time PCR genotyping assay.

The TaqMan real-time PCR genotyping assay established in this study can be considered a quicker method with higher throughput to perform genotyping for *PKD1* when compared to the pre-existing methods. Sanger sequencing is the most traditional albeit concrete method for decoding sequences and is still used for confirmation because of its high accuracy. While it is important to refer to Sanger sequencing when establishing a new genotyping method, there is no need to read the entire sequence when detecting SNPs, and easier methods can be used for genotyping. Recently, the PCR-restriction fragment length polymorphism method has been used widely for SNP detection because of its reproducibility. However, both these methods involve many steps, which could take several hours. On the other hand, our genotyping method can be run on a fast setting on real-time PCR instruments and takes only up to 40 min. Additionally, this method allows the analysis of 93 samples at once (on a 96 well-plate, with one negative control and two positive controls), making genotyping of multiple samples remarkably faster and more convenient. TaqMan real-time PCR genotyping method has sufficiently high specificity and is compatible with other methods, making it a very useful and potential substitute for pre-existing methods.

According to the real-time PCR genotyping assay, the conventional *PKD1* variant was observed in 1.8% of all included cases, which was relatively lower than that in previous reports. The prevalence of the variant was 13.5% in China (n=111), 33.3% in Slovenia (n=24), and most surprisingly, 40% in the study conducted in 2019 in Japan (n=377) [11, 18, 25]. The huge difference between the results of our study and the 2019 Japanese study is most likely due to the difference in the research methods; the previous study focused on cats already suspected of ADPKD based on imaging or family history, as opposed to our study, which included cats regardless of their medical history. The previous study can be considered more inclined toward relationship analysis between ultrasonographic and genetic test results rather than an epidemiological study. Thus, the overall prevalence of the variant in Japanese cats is presumably closer to the 1.8% obtained in this study. Similarly, the study in Slovenia was limited to Persian cats, which could explain the high prevalence reported. The method used in the study from China was closest to ours; however, the number of cases included was much less, which may have increased the reported prevalence of the variant. However, this could also be a reflection of the regional differences.

The odds of having the conventional *PKD1* variant were significantly higher in Persian cats and Exotic Shorthairs, a cross between Persians, and the short-haired breeds such as American or British Shorthairs, corroborating the findings of previous studies [2, 3, 8, 12, 18]. Scottish Folds also had a significantly high odds ratio, similar to that reported in the previous research conducted in Japan [25]. Although some American Shorthairs had the conventional *PKD1* variant, their odds ratios were not significant due to their large population. Furthermore, 6 out of 718 mixed breed cats presented the variant, suggesting that mixed breeds are also potential carriers of the conventional *PKD1* variant in Japan.

In cats with the conventional *PKD1* variant, the BUN levels were significantly higher in those with renal cysts than in those without, suggesting that the presence of renal cysts is associated with renal function deterioration. This finding supports the notion that renal function deteriorates in ADPKD. However, there was no significant difference in the plasma creatinine levels of patients with and without renal cysts, though they tended to be higher in the former than in the latter (*P*=0.0989). This could be due to the small number of ADPKD cases in this study, and further research regarding renal function in ADPKD cases in a larger population is warranted. We could not evaluate the relationship between the cyst size or number and the deterioration of renal function, and determining whether the increase in cysts actually correlates with renal function decline is important for developing effective treatment methods. Future research is needed to clarify this, ideally over the course of a few years, in the same individuals. The results of the real-time PCR *PKD1* genotyping assay mostly reflected the results of the ultrasonographic examination, with a few exceptions. Four cats with the conventional *PKD1* variant did not have renal cysts, which could be attributed to the stronger tendency for cysts to be present in older cats than in younger cats. Notably, these four cats were aged 9 months, 11 months, 5 years, and 8 years, respectively; the two older cats had cysts in the liver and pancreas, respectively, which are common complications of ADPKD. Interestingly, there were nine negative cases with cysts in the kidneys, liver, or pancreas, suggestive of ADPKD. This suggests the possibility of a different variant causing the disease, as indicated in previous studies [13]. Since human ADPKD, a disease very similar in pathology and mechanism to feline ADPKD, is caused by many different variants of *PKD1* and even some other genes involved in transportation of PC1 protein [22], it is only natural that feline ADPKD may have other causative variants too.

To explore unknown *PKD1* variants that could be the additional cause of ADPKD, we performed target resequencing using next-generation sequencing. As a result, 86 novel variants were detected in total, and when we focused on ADPKD cases without the conventional *PKD1* variant (chrE3:g.42858112C>A), 27 variants were extracted. Of these variants, four singleton variants were found only in cases with ADPKD and in none of the unaffected cats, indicating their association with the disease. All four variants were in exon 15, suggesting that along with exon 29, exon 15 also plays an important role in the cystogenesis of ADPKD. ChrE3:g.42848725delC was the only variant that caused a frameshift, and although its impact could not be calculated using prediction tools such as PolyPhen-2, it would definitely damage the PC1 structure and function substantially. Of the remaining three missense variants, chrE3:g.42850283C>T was the only one to be probably damaging as predicted by PolyPhen-2. Interestingly, when we compared the location of human genes using the UCSC LiftOver tool, this variant was the only variant that had a complete match with a known variant of the human *PKD1*. This suggests that the same variant can lead to ADPKD in both humans and cats, thus taking the resemblance between feline and human ADPKD and the role of feline ADPKD as a pathological model to another level.

Our study was based on patients referred to the University of Tokyo Veterinary Medical Center, which specializes in secondary care. Thus, it may have had more number of ill animals than would a primary care clinic or the Japanese animal population as a whole. We focused on variants that were present only in ADPKD cases and did not analyze variants that were found in both ADPKD cases and a few control cases further. As shown in Table 4, several variants from ADPKD cases were also seen in less than five control cases. We cannot deny the possibility of those cases being misdiagnosed as “healthy” or simply being too young to develop detectable cysts, as most of the samples were scanned by ultrasonographic examination, meaning that there may be overlooked variants that actually were the cause of ADPKD. To rule out this possibility, we need to monitor these “control cases” carefully to determine if they will develop ADPKD in the future. The four new variants proposed in this study were only found once each in three out of six ADPKD samples lacking the nonsense transversion on chrE3:g.42858112C>A. In future research, the collection of more ADPKD samples lacking the nonsense variant is required to study the frequency of the four variants. Additionally, although two of these variants were predicted to be benign, considering that they were found in cats with ADPKD, there is a possibility that these variants have a destructive effect on the resulting protein. Hence, protein analysis is necessary to evaluate the impact of these variants correctly.

In summary, the conventional *PKD1* variant (chrE3:g.42858112C>A) was found to be most common in Scottish Folds, Persian cats, and Exotic Shorthairs in Japan, and four new variants were suggested as the cause of feline ADPKD. These new variants were all found in exon 15 of *PKD1*, indicating that exon 15 along with exon 29 are key factors in ADPKD pathogenesis. By promoting further *PKD1* analysis, fundamental prevention of the disease can be achieved by controlling the breeding of *PKD1* variant carriers.

## CONFLICT OF INTEREST

The authors declare that there is no conflict of interest.

## ACKNOWLEDGEMENTS

We would like to thank Drs. M. Yoshida and M. Sakaguchi (Azabu University) for providing the feline DNA samples and Dr. M. Arahori and Ms. K. Koinuma for helping with target sequencing. This study was supported by JSPS KAKENHI, a Grant-in-Aid for Science Research (Grant Number:19H00968), and the Anicom Capital Research Grant (EVOLVE).

## Notes

### Competing Interest Statement

The authors have declared no competing interest.

